# Reactivation of reward-related patterns from single past episodes supports memory-based decision making

**DOI:** 10.1101/035196

**Authors:** G. Elliott Wimmer, Christian Büchel

## Abstract

Rewarding experiences exert a strong influence on later decision making. While decades of neuroscience research have shown how reinforcement gradually shapes preferences, decisions are often influenced by single past experiences. Surprisingly, relatively little is known about the influence of single learning episodes. While recent work has proposed a role for episodes in decision making, it is largely unknown whether and how episodic experiences contribute to value-based decision making and how the values of single episodes are represented in the brain. In multiple behavioral experiments and an fMRI experiment, we tested whether and how rewarding episodes could support later decision making. Participants experienced episodes of high reward or low reward in conjunction with incidental, trial-unique neutral pictures. In a surprise test phase, we found that participants could indeed remember the associated level of reward, as evidenced by accurate source memory for value and preferences to re-engage with rewarded objects. Further, in a separate experiment, we found that high reward objects shown as primes before a gambling task increased financial risk-taking. Neurally, re-exposure to objects in the test phase led to significant reactivation of reward-related patterns. Importantly, individual variability in the strength of reactivation predicted value memory performance. Further, local searchlight analyses identified significant reactivation in the ventromedial PFC. Our results provide a novel demonstration that affect-related neural patterns are reactivated during later experience. Reactivation of value information represents a mechanism by which memory can guide decision making.

## Introduction

Value-based decision making is an essential component of adaptive behavior. Such decisions can be based on different types of knowledge: When choosing a restaurant, we could inform our choice using a life-long preference for a favorite food or instead by a single positive memory of a new type of cuisine. Learning and decision making research has largely focused on the former, studying well-learned values built up over many repeated experiences (Daw and Doya, 2006; Schultz, 2006; Rangel et al., 2008). Often, however, we make decisions based on information provided by specific past experiences. Surprisingly, it is largely unknown how episodic experiences contribute to value-based decision making. As strongly reinforcing experiences are rare, limiting the utility of gradual learning mechanisms, it may be particularly important to remember the value of single episodes.

Decision making and episodic memory are often studied separately. However, value-based choice in some contexts has been proposed to be supported by a mechanism that samples representations stored in memory (Weber and Johnson, 2006; Biele et al., 2009; Gluth et al., 2015). In these proposals, it is necessary that episodes are encoded along with their value. Providing some support for this possibility, behavioral experiments have shown that in a situation where participants can compare the value of two past episodes (e.g., different experiences of unpleasant cold water) (Kahneman et al., 1993; Redelmeier and Kahneman, 1996). However, it remains unknown how the value of single episodes may be represented in the brain.

From the perspective of memory, a rich literature has studied the modulation of memory by emotion, a concept that may be closely related to value and reward. This research has shown that both positive and negative emotion often increase memory strength (for review, see Reisberg and Heuer, 2004; LaBar and Cabeza, 2006). Interestingly, it has been shown that reward value can incidentally facilitate memory (Wittmann et al., 2005; Mather and Schoeke, 2011; Gruber et al., 2014; Murayama and Kitagami, 2014; Murty and Adcock, 2014; Koster et al., 2015). In these studies, incidental information shown during high vs. low reward anticipation or curiosity is remembered better, and this benefit is reflected in striatal, midbrain, and hippocampal activity (Wittmann et al., 2005; Gruber et al., 2014). However, whether people automatically and implicitly encode the value associated with single experiences, as reflected in later source memory and decision making behavior, has not been a focus of memory research. Successful incidental encoding of the affective value of previous episodes could allow later reactivation of value associations to guide decision making.

Critically, it remains unknown whether affect-related patterns of brain activity from encoding are present during the reactivation of memories. Researchers have utilized designs where neutral items are associated with emotional contexts during encoding and then the neutral items are re-presented at test (for review, see Buchanan, 2007), thus avoiding problems associated with re-presenting intrinsically emotional stimuli. With univariate analysis methods, imaging results suggest some activation of affect-related brain regions during re-exposure, based on overlap of mean group effects or inferential reasoning from other studies (Buchanan, 2007; Tsukiura and Cabeza, 2008). More recently developed multivariate analysis methods, which provide improved specificity by detecting distributed patterns of brain activity, offer a promising approach for answering this question (Kriegeskorte et al., 2006; Rissman and Wagner, 2012).

In the following experiments, we investigated whether source memory for episodic value associations and value-based decision making could be supported by and influenced by single rewarding episodes. Our experiments examined episodic memory in terms of single-experience incidental encoding of events in conjunction with reward. In a motivated reward task, neutral objects were presented once, incidentally paired with high or low reward. A surprise test for the value associated with the objects followed, allowing us to measure whether the value episodically associated with an item influenced source memory and decision making. Importantly, in order to match real-world learning situations where agents are not aware in advance of what information may be useful to remember, the encoding of reward-episode associations in our experiments was implicit.

## Materials and methods

**Participants**. A total of 92 subjects participated in Experiments 1-4 and separate set of 30 subjects participated in the fMRI study. Participants were fluent German speakers with no self-reported neurological or psychiatric disorders and normal or corrected-to-normal vision. In Experiment 1 on reward value memory, 31 subjects were included (25 female; mean age, 25 years; range, 19-34 years). In Experiment 2 on reward vs. pain value memory, data from 1 subject were excluded due to thermode failure, leaving 20 subjects (13 female; mean age, 26 years; range, 20-31 years). In Experiment 3 on reward episode re-engagement, 20 subjects were included (10 female; mean age 24 years; range 18-30 years). In Experiment 4 on object-primed risk taking, data from 1 subject were excluded due to insufficient high-risk choices (less than 15% of trials), leaving 20 subjects (12 female; mean age 24 years; range 19-28 years). In the fMRI study, data from 1 subject were excluded due to a misunderstanding of instructions, leaving 29 right-handed subjects (16 female; mean age, 25 years; range, 19-30 years). All subjects were remunerated for their participation, and participants were additionally compensated based on choice performance or winnings. The Ethics committee of the Medical Chamber Hamburg approved the study and all subjects gave written consent.

**Behavioral and fMRI procedure: Incidental learning phase**. The reward value memory task consisted of an incidental learning phase and a value memory test phase. The incidental learning phase exposed participants to incidental trial-unique objects during the experience of high or low reward anticipation and outcome to allow for the incidental encoding of episodes associated with high or low reward value (Fig. 1*A*). Color pictures of objects were drawn from a previously used set of images compiled via internet search (Wimmer et al., 2014), largely composed of familiar non-food household items set on white backgrounds. We used an adapted Monetary Incentive Delay (MID) task (Knutson et al., 2000; Mather and Schoeke, 2011). Participants were instructed that one of two reward amounts would be available on a trial, a high reward (€2.00) or low reward (€0.01). Critically, to prevent the use of visual association memory to guide responses in the later value memory test, the reward cue and reward feedback did not use text or pictorial information to indicate reward. Instead, reward was indicated by the orientation of object shading, and this shading reversed at the mid-point of the task, accompanied by an extensive retraining phase preceding trials of interest. Importantly, the incidental nature of learning made the use of explicit encoding strategies very unlikely.

**Figure 1.**
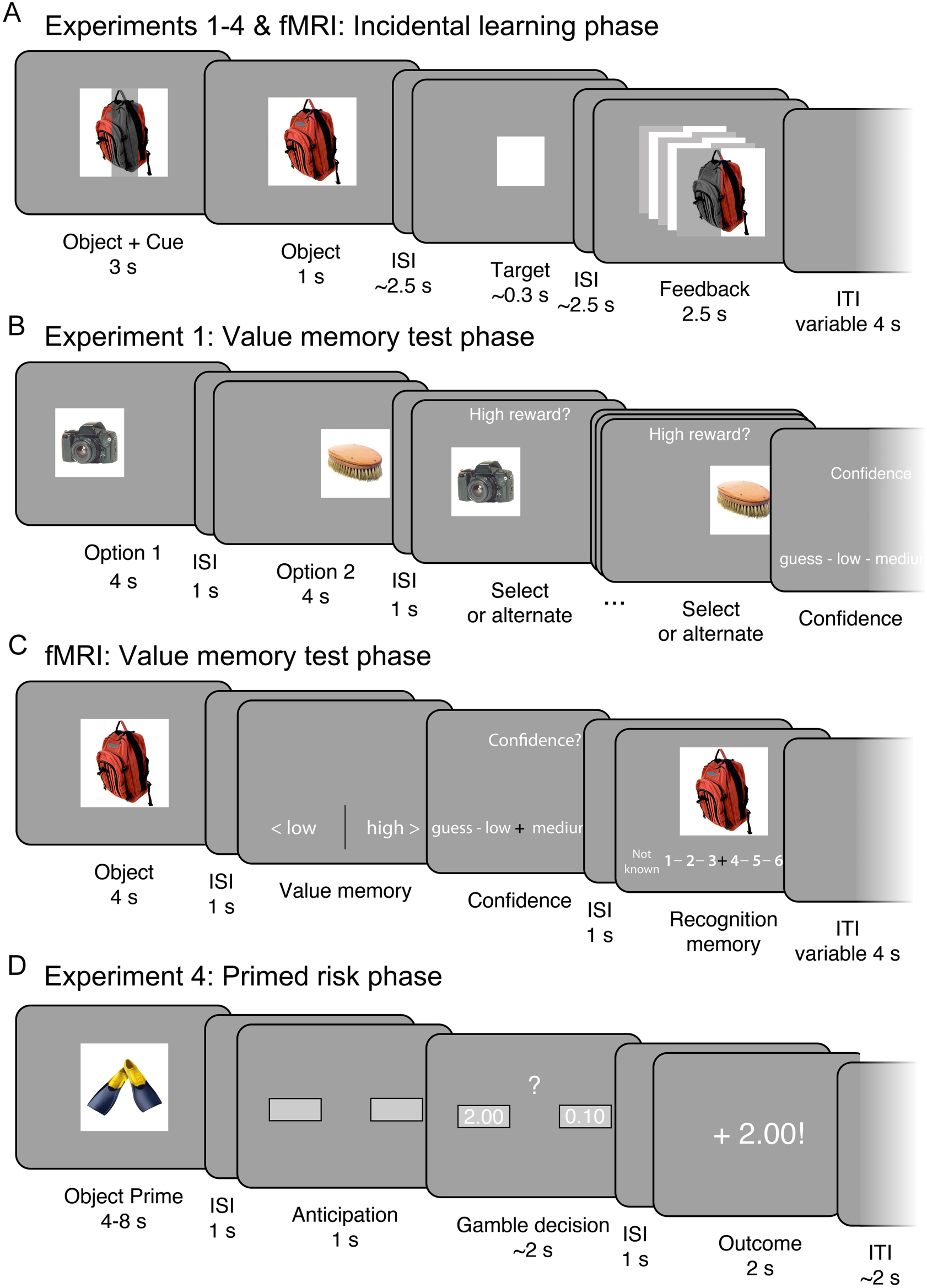
Reward value memory experiment. ***A***, Reward cue value (€2.00 or €0.01) was indicated by a shaded bar across the incidental object picture, after which participants made a rapid response to the target. Reward feedback was indicated by flickering halves of the object (cue and feedback shading was counterbalanced across blocks). ***B***, In Experiment 1, a surprise source memory choice test phase followed the incidental learning phase. Participants were incentivized to choose the object associated with higher reward in the incidental learning phase. In Experiment 2, participants were incentivized to choose the object associated with a “Better experience” (reward vs. pain). In Experiment 3, participants chose which object they would like to re-engage with (“Play again”) in an additional reward phase. ***C***, In the fMRI experiment, a surprise value memory test phase followed in which participants responded with whether objects had been associated with high vs. low reward. ***D***, In Experiment 4, high and low reward-associated objects were presented prior to an unrelated gamble decision to examine whether reward episodes exert an influence on unrelated value-based decisions.

On each trial an incidental object was presented for 3 s. For the first 2 s, the object was shown with a vertical or horizontal shaded bar signaling the potential for high or low reward. After a variable ISI (1.5 s minimum, plus a value randomly drawn from a uniform 0-2 s distribution), the white square target appeared to which participants were instructed to rapidly press a button. The target duration was adjusted for each subject throughout the task to achieve an approximately 70% hit rate. A variable ISI (1.5 s minimum, plus the difference between 2 s and the pre-target ISI random value) followed the target. Then, feedback was presented for 2.5 s in conjunction a re-presentation of the object picture from the cue. If participants responded in time to the target, they received the reward indicated at the start of the trial; if the response was too slow, participants received no reward. Reward feedback was indicated by a vertical or horizontal flickering of the two halves of the object picture, congruent with the cue. For example, if the reward cue was a vertical bar, a successful trial would indicate reward feedback via alternating flickering of the left and right side of the object picture (Fig. 1*A*). Miss trials presented the object in grey-scale with no flicker. Trials were followed by a variable 4 s mean (range: 2-10 s) inter-trial-interval. Two pseudo-random orderings of incidental object pictures were used for counterbalancing object and reward associations.

Before entering the MRI scanner, participants completed 16 practice trials, with an additional 4 practice trials completed inside the MRI before the start of the experiment. The practice trials allowed for learning of the high-value horizontal shaded bar and low-value vertical shaded bar cue-reward associations. Practice was also used to calibrate the initial target duration. During practice trials, the potential value of the trial was additionally indicated in text on the left and right side of a centrally presented, trial-unique, abstract color stimulus. Reward feedback was also indicated in text on the left and right side of the picture. In the actual experiment, no text reward information was presented.

After an initial block of 34 trials, participants were instructed that the vertical and horizontal reward cues reversed. Following this instruction, to establish learning of the new cue contingencies, participants completed an additional 20 practice trials, where trial-unique abstract color stimuli and flanking text reward information was included. After this re-learning practice phase, a second experimental block of 34 trials was presented where no text reward information was presented. In the two incidental learning phase blocks, a total of 34 high reward trials and 34 low reward trials were presented. In each block, after 17 trials a pause of 30s was inserted to allow for a brief rest period.

Results from the incidental learning phase demonstrate that reaction times were faster for the high vs. low reward cue, and that the hit rate titration procedure was successful. In behavioral Experiment 1, we observed a high reward hit rate of 69.2 ± 1.0% and a low reward hit rate of 70.5 ± 1.1% (mean ± SEM; high-low difference, p = 0.38). Reaction times were faster for the high reward target than the low reward target (high reward RT: 391.5 ± 19.1 ms; low reward RT: 431.3 ± 18.1 ms; t_(28)_ = 2.23, p = 0.033), suggesting that the reward manipulation successfully affected behavior. In Experiment 2, we observed a high reward hit rate of 69.6 ± 1.2% and a low reward hit rate of 68.2 ± 1.6% (high-low difference, p = 0.50). Reaction times were faster for the high reward target than the low reward target (high reward RT: 294.3 ± 10.2 ms; low reward RT: 311.9 ± 6.8 ms; t_(19)_ = 2.40, p = 0.027). In Experiment 3, we observed a high reward hit rate of 72.2 ± 1.0% and a low reward hit rate of 71.4 ± 1.2% (high-low difference, p = 0.61). Reaction times were faster for the high reward target than the low reward target (high reward RT: 305.5 ± 12.3 ms; low reward RT: 325.7 ± 7.8 ms; t_(19)_ = 2.84, p = 0.01). In Experiment 4, we observed a high reward hit rate of 70.4 ± 1.1% and a low reward hit rate of 68.0 ± 1.0% (high-low difference, t_(19)_ = 2.48, p = 0.023); note that this difference in hit rate does not affect our ability to examine the effect of reward episodes on decision making. Reaction times tended to be faster for the high reward target than the low reward target (high reward RT: 317.0 ± 13.7 ms; low reward RT: 331.7 ± 13.8 ms; t_(19)_ = 1.93, p = 0.069). In the fMRI experiment, we observed a high reward hit rate of 70.3 ± 1.1% and a low reward hit rate of 69.0 ± 0.9% (high-low difference, p = 0.42). As in the behavioral experiments, reaction times were faster for the high reward target than the low reward target (high reward RT: 249.0 ± 6.6 ms; low reward RT: 259.4 ± 7.9 ms; t_(28)_ = 2.49, p = 0.019).

**Behavioral procedure: Experiment 2 pain incidental learning phase**. In Experiment 2, to control for the effect of arousal on source memory judgments for value, we compared incidental reward-associated objects to pain-associated objects. We collected one half of the above-described reward incidental learning phase and a separate pain incidental learning phase where objects were associated with thermal heat pain. The order of the reward incidental learning phase and the heat calibration and pain incidental learning phases were counterbalanced across participants.

Before the pain incidental learning phase, heat levels were calibrated for each participant to achieve the same subjective high and low aversive pain experience across participants. Thermal stimulation was delivered via an MRI compatible 3 × 3 cm Peltier thermode (MSA; Somedic, Sweden), applied to the inner left forearm. During the visual presentation of a white square, heat was applied for 10 s. For pain rating, we used a 1-8 rating scale with 0.5-point increments, superimposed on a yellow-to-red gradient. An arrow cursor was moved from the initial mid-point starting location using left and right key-presses and ratings were confirmed with the space bar. A rating of ‘8’ corresponded to the highest level of heat pain a subject could endure multiple times. If the level of pain was intolerable, participants moved the rating past the ‘8’ end of the scale, at which point a ‘9’ appeared on the screen. Participants rated the pain associated with a pseudorandom list of 10 different temperatures ranging from 39.5 to 49.5°C. A linear interpolation algorithm then selected a low temperature estimated to yield a ‘2’ rating and a high temperature estimated to yield a ‘7.5’ rating.

Next, across 2 mini-blocks, 17 high heat trials and 17 low heat trials were presented. On each trial, the incidental object appeared in conjunction with thermal heat. To allow for a better match between the appearance of the object and the onset of noticeable heat, heat onset started 0.75 s prior to object appearance (for a similar method, see Forkmann et al., 2013). The incidental object was presented for a total duration of 10 s, after which the temperature returned to baseline (33°C) over several seconds. After a 4 s ISI, the pain rating scale appeared. Participants used the left and right buttons to move a selection arrow from the initial cursor position (randomized between 4.5-5.5) to their experienced pain level and pressed the down button twice to make their selection; responses were self-paced. After the participant entered their response, trials were followed by a variable 2 s mean (range: 0.5-6 s) inter-trial-interval (ITI). To maintain attention on the screen during object presentation, on a random 50% of trials the object picture flickered in illumination during heat stimulation. When a flicker was detected, participants were instructed to press the down button. During the break between the two incidental learning phase blocks, the thermode was moved to a new location on the inner arm to avoid sensitization.

To maintain similar differences in subjective experience between the high and low heat conditions, temperatures were automatically adjusted throughout the task to maintain the targeted pain rating values. If the median of the previous 6 low heat trials fell below a rating of 1.5, the low temperature was increased by 0.2°C; if the median rating was above 3, the low temperature was decreased by 0.2°C. For the high temperature, if the median rating fell below 7.5, the high temperature was increased by 0.2°C (if the temperature was below 50.5°C). If a rating of “9” was given, indicating an intolerably high level of pain, the high temperature was decreased by 0.8°C. Overall, in Experiment 2 the ratings for the high and low heat reliably differed (low pain rating, 2.4 ± 0.2; high pain rating, 7.3 ± 0.10). The mean administered low temperature was 41.1 ± 0.4°C and the high temperature was 49.7 ± 0.3°C.

**Behavioral procedure: test phase**. In Experiment 1, we administered a surprise memory test for the value of the single episodic experiences in a test phase (Fig. 1*B*). We used an incentivized choice task to examine source memory for episodic value associations. Participants’ goal was to choose which of two incidental objects had been presented during high reward in order to earn bonus monetary rewards. Correct choices of the high reward option were rewarded with €0.50, given after the experiment (no trial-by-trial feedback was presented). The presentation of the choice options was separated in time to allow for better recall of the value of individual objects. From a memory perspective, choice in this task is a source memory judgment.

On each of the 34 choice trials, first, one object, randomly selected, was presented randomly in the left or the right half of the screen for 4 s. After a 1 s ISI, the other image was presented in the other half of the screen. After a 1 s ISI, one of the items, randomly selected, re-appeared in the position it had occupied initially.

Participants could select the currently displayed object with the space key or view the other object using the left or right arrow keys. For example, if the subject wished to view the other object and the current object occupied the left of the screen, the right arrow would reveal the other object. Participants could move between the two objects as many times as they wanted until finding the object remembered to be associated with high reward. Next, participants rated the confidence in their choice on a 1-4 “not at all” to “very” confident scale. Choice trials were followed by a variable 3 s mean (1-10 s range) ITI.

The individual choice options for the choice trials were selected using an algorithm that paired high and low reward-associated objects from successful hit trials (where both the €2.00 and €0.01 amounts were won), or alternatively between high and low reward-associated objects from unsuccessful miss trials. Any remaining objects after this procedure were paired in mixed hit and miss choices. To eliminate the possibility of reward cue orientation information (the horizontal or vertical shading) influencing choice, both objects in a choice were selected such that they had been associated with the same cue orientation. This was accomplished by selecting objects from different halves of the task. For example, a high reward-associated object would be taken from the first half of the task and a low reward-associated object would be taken from the second half of the task. This algorithm was individualized for each participant and the order of choices was randomized. A practice trial preceded the start of the choice phase to familiarize participants with the task; the practice choice options were a picture of a euro coin and a red circle with €0.00 indicated.

In Experiment 2, we tested whether general arousal differences between the high and low reward-associated objects could account for test phase performance by comparing reward-associated vs. pain-associated experiences. We employed a similar test phase design as in Experiment 1, with the exception that choices were between objects that had been incidentally paired with reward vs. objects that had been incidentally paired with thermal heat pain. The choice prompt within the task, instead of “Higher reward?” as in Experiment 1, was “Positive experience?”. In 14 critical choice trials, participants chose between a high reward-associated object and a high pain-associated object. In the remaining choice trials (not analyzed for the present report), choices were either within-valence (e.g. high reward vs. low reward) or across-valence (low reward vs. low pain).

In Experiment 3, we tested whether participants could also make future-guided decisions about the reward-associated objects without reference to recalling their previous associations in the incidental learning phase. We employed a similar test phase design as in Experiment 1, with the critical exception of the instructions for the choice between objects. Participants were told that their goal was to select objects to “re-play” in another round of the fast response game (the adapted MID task in the incidental learning phase). They were told to try to select the picture that they thought would be “luckier” and which would win them more money in another round of the game. The choice prompt within the task, instead of “Higher reward?” as in Experiment 1, was “Play again?”. After this block, participants were rewarded based on the number of high-reward objects they selected in the test phase (in place of actually re-playing the learning phase task).

**Behavioral procedure: Experiment 4 reward object-primed risk taking**. In Experiment 4, we utilized a risk-taking gambling task to examine whether objects that had been incidentally paired with reward also influenced unrelated value-based decisions (Fig. 1*D*). Our design was adapted from Knutson et al. (2008), who used a similar experimental design, but with emotional IAPS pictures acting as pre-gamble “primes”. The incidental learning phase was the same as in Experiments 1-3 with the exception that the low reward amount was €0.10 instead of €0.01.

In the risk taking task, participants were endowed with €10 to start the game, to which their gambling results were added or subtracted. Before each gamble decision, participants first attended to the presentation of a gamble-unrelated object. Although participants were not given this information in the instructions, objects were drawn from the preceding incidental reward phase. After the object prime, a gamble decision appeared with a €2.00 high bet option and a €0.10 low bet option. While the expected values of the two options were both €0.00, the high bet option had a larger variance than the low bet option, and thus we will refer to the high bet option as the “risky” option and choices for the high bet option as “risky” choices (Knutson et al., 2008). Participants were instructed that they would win around half of the time and lose about half of the time for a given gamble decision, and that for each choice, they should go with their best feeling about which option would be better at that point in the game

On each gamble trial, an object prime was initially presented for 4-8 s (Fig. 1*D*). Participants’ goal was to press the space bar if and when the object flickered in brightness. Objects flickered 0, 1, or 2 times for 0.35 s, and the total duration of object presentation was 4 s, 6 s, or 8 s, respectively, for the different number of flickers. A 1 s ISI followed. Next, the unrelated gamble decision phase began. Empty response option boxes were first presented during a 1 s anticipation phase, after which the high and low gamble amounts were displayed. The left/right presentation of the gamble options was randomized. Participants had 4 s in which to make their gamble decision. A 1 s ISI immediately followed the response. The gamble outcome (reward or miss) was then presented for 2 s. To increase interest in and attention toward the game, the high gamble outcome amounts varied in a normal distribution around the ±2.00 outcome with a €0.125 standard deviation (e.g. €2.12); for the low risk gamble, amounts varied in a flat distribution around the ±0.10 outcome with a range of ±0.05 (e.g. 0.09). The task was divided into 6 mini-blocks, and in the trial following a block break (as well as the first trial) a novel object picture preceded the gamble; these trials were excluded from analysis. During the breaks, participants were informed whether their current total winnings were above the initial €10 endowment or below €10. A practice trial preceded the start of the gamble phase to familiarize participants with the task.

The number of “flickers” for an object was drawn from a pseudorandom list, with flicker amount balanced across high and low reward object prime trials. Win vs. miss gamble outcome was drawn from one of four pseudorandom lists, with feedback balanced across high and low reward object prime trials. For the gamble choice, to ensure sufficient numbers of risky and non-risky choices (as we had no interest in absolute risk preferences), the high risk option amount was increased if participants exhibited excessive risk-taking or decreased if participants exhibited infrequent risk-taking. Specifically, after every 12 critical object-primed trials in the first 4 mini-blocks of the task, if a participant had chosen the high gamble on more than 9 out of 12 preceding trials, the high risk option was increased by €1.00 (up to €3.00 maximum). Conversely, if a participant had chosen the low gamble on more than 9 out of 12 preceding trials, the high risk option was decreased by €1.00 (or if the current value was €1.00, the high risk option was decreased to €0.50). Participants were notified of any change in gamble amounts before the start of the next mini-block. However, most participants showed a balance of choices of the high and low gamble amounts (mean risk-taking rate, 50.2 ± 2.5%; min = 27.7%, max = 71.2%). Thus, the high-risk option frequently remained at the initial €2.00 value (mean, €1.98 ± 0.09). Further, control multilevel regression analyses did not find any interaction between the high gamble amount and the influence of object primes.

**fMRI Procedure: test phase**. After the scanned incidental learning phase, a high-resolution structural scan was acquired. To test memory for the incidental episodic reward associations, participants completed the surprise reward value memory test (Fig. 1*C*).

On each trial a single object was presented alone for 4 s to allow participants to remember the reward association in the absence of response demands. Next, after a 1 s ISI, participants were presented with the value memory question, with response options of “low reward” and “high reward”. Participants pressed the left or right buttons to indicate whether they thought that the object had been associated with low reward or high reward cue information in the incidental learning phase. To minimize response bias, participants were instructed that half of the objects had been incidentally shown during high reward information trials and half with low reward information in the preceding phase. All test phase responses were self-paced. Next, after a 0.5 s ISI, a confidence rating screen appeared with 4 levels of response: “guess”, “somewhat certain”, “certain”, and “very certain”. Participants used the left and right buttons to move from the initially presented center point (between the middle two options) to their selected response and then pressed the down button to make their selection. Finally, after a 1 s ISI, a 6-point memory recognition strength scale was presented. Participants indicated by selecting a number on the scale to what degree they thought the object was “not known” or “definitely known”. Participants used the left and right buttons to move from the initially presented center point (between the “guess” responses) to their selected response and then pressed the down button to make their selection. A variable ITI with a mean of 4 s (range: 2-10 s) followed. The order of the old pictures was pseudo-randomized from the incidental learning phase order. The memory recognition scale was reversed in half of the participants.

For the incidental learning and value memory test phases of the reward experiments, the duration and distribution of ITIs (or “null events”) was optimized for estimation of rapid event-related fMRI responses as calculated using Optseq software (http://surfer.nmr.mgh.harvard.edu/optseq/). The experimental tasks were presented using Matlab (Mathworks, Natick, MA) and the Psychophysics Toolbox (Brainard, 1997). The behavioral experiment was completed on a laptop computer. In the fMRI experiment, the task was projected onto a mirror above the subject’s eyes. Responses were made using a 4-button interface with a “diamond” arrangement of buttons (Current Designs, Philadelphia, PA). At the end of the experiment, participants completed a paper questionnaire querying their knowledge of the task instructions and their expectations (if any) regarding the incidental object pictures. In response to a question about whether the fMRI participants expected to be tested in some way about the objects (including simple memory for the pictures), 43.1 ± 8.9% answered “yes”. However, we found no difference in behavioral performance between those who expected some kind of memory test and those who did not (p = 0.92). Finally, task instructions and on-screen text were presented in German; for the methods description and task figures, this text has been translated into English.

**fMRI Data Acquisition**. Whole-brain imaging was conducted on a Siemens Trio 3 Tesla system equipped with a 32-channel head coil (Siemens, Erlangen, Germany). Functional images were collected using a multiband acquisition sequence (TR = 1240 ms, TE = 26 ms, flip angle = 60, multiband factor = 2; 2 × 2 × 2 mm voxel size; 40 axial slices with a 1 mm gap). Slices were tilted approximately 30° relative to the AC–PC line to improve signal-to-noise ratio in the orbitofrontal cortex (Deichmann et al., 2003). Head padding was used to minimize head motion. No subject’s motion exceeded 3 mm in any direction from one volume acquisition to the next, and in the single case of 3 mm motion, this volume and surrounding volumes were replaced by the mean of adjacent TRs. For each functional scanning run, four discarded volumes were collected prior to the first trial to allow for magnetic field equilibration.

During the incidental learning phase, two functional runs of an average of 438 TRs (9 min and 3 s) were collected, each including 34 trials. During the memory test phase, three functional runs of an average of 294 TRs (6 min and 4 s) were collected, including 22-23 trials. For one subject in the test phase, fMRI data acquisition failed during the second test run; for this subject, fMRI analysis included runs 1 and 3.

Structural images were collected using a high-resolution T1-weighted magnetization prepared rapid acquisition gradient echo (MPRAGE) pulse sequence (1 × 1 × 1 mm voxel size) between the incidental learning phase and the value memory test phase.

**Behavioral Analysis**. The primary behavioral question was whether participants exhibited memory for the value associated with the incidental object pictures. Multilevel regression models as implemented in R (R-project.org) were used to further investigate value memory and recognition memory. All models entered subject as a random effect. Additionally, a simple reinforcement learning model was also used to generate predictive trial value variables of interest. This model used a Rescorla-Wagner update rule (Sutton and Barto, 1998), where a high reward trial set to a reward value of 1 and a low reward trial set to a reward value of zero, with a learning rate of 0.5. As hit vs. miss feedback had no effect on value memory behavior (see Results), we did not expect reward learning models based on feedback success; indeed, alternative learning models based on feedback showed no significant effects. In Experiment 4, control multilevel regression analyses confirmed that object prime duration and the interaction of object value and duration did not affect risk-taking and switches to the risky option.

**fMRI Data Analysis**. Preprocessing and data analysis was performed using Statistical Parametric Mapping software (SPM8; Wellcome Department of Imaging Neuroscience, Institute of Neurology, London, UK). Before preprocessing, individual slices with artifacts were replaced with the mean of the two surrounding timepoints using a script adapted from the ArtRepair toolbox (Mazaika et al., 2009). Images were realigned to correct for subject motion and then spatially normalized to the Montreal Neurological Institute (MNI) coordinate space by estimating a warping to template space from each subject’s anatomical image and applying the resulting transformation to the EPIs. Images were filtered with a 128 s high-pass filter and resampled to 2 mm cubic voxels. For univariate analyses, images were then smoothed with a 6 mm FWHM Gaussian kernel.

fMRI model regressors were convolved with the canonical hemodynamic response function and entered into a general linear model (GLM) of each subject’s fMRI data. The six scan-to-scan motion parameters produced during realignment were included as additional regressors in the GLM to account for residual effects of subject movement.

We first conducted “localizer” univariate analyses to identify main effects of reward in the incidental learning phase of the fMRI studies. The GLM included regressors for the cue onset (3 s duration), target (0 s duration), and feedback (2.5 s duration). The cue regressor was accompanied by a modulatory regressor for high vs. low reward and the feedback regressor was accompanied by a modulatory regressor for hit vs. miss feedback.

We also conducted exploratory univariate analyses to examine the modulation of brain activity by the test phase re-presentation of object pictures. This model included regressors during the object re-presentation (4 s), value memory response (variable duration), confidence rating (variable duration), and recognition memory rating (variable duration). The object re-presentation regressor was modulated by 3 variables: the reward incidentally associated with the object during the learning phase, the confidence rating given on that trial, and the memory response given on that trial. We conducted an additional control analysis where the memory response was entered first in the GLM.

**Regions of interest**. For multivariate classification analyses, we constructed a mask from voxels responsive to high vs. low reward anticipation in the “localizer” GLM described above at an uncorrected threshold of p < 0.001 (excluding the cerebellum) and voxels responsive to hit vs. miss feedback in a contiguous cluster including the ventromedial prefrontal cortex (VMPFC), striatum, and hippocampus (excluding the cerebellum and any posterior regions that may show activation purely based on the flicker-related perceptual difference between hit and miss trials) at an uncorrected threshold of p < 0.001. Note that different thresholds for the ROI mask yielded qualitatively similar results.

For univariate and searchlight results, linear contrasts of univariate SPMs were taken to a group-level (random-effects) analysis. We report results corrected for family-wise error (FWE) due to multiple comparisons (Friston et al., 1993). We conduct this correction at the peak level within small volume ROIs for which we had an a priori hypothesis (after an initial thresholding of p < 0.005 uncorrected) or at the whole-brain cluster level, with a cluster threshold of 10 voxels. We focused on reward-related responses supporting memory for value in the striatum and VMPFC. The striatum ROI was adapted from the AAL atlas (Tzourio-Mazoyer et al., 2002). As the VMPFC is known to correlate with reward value as well as to respond to reward feedback in the MID task (Knutson et al., 2001; Bartra et al., 2013; Clithero and Rangel, 2014), we constructed the VMPFC ROI from the contrast of hit vs. miss feedback in the incidental learning phase, thresholded at p < 0.00001 uncorrected. The striatum and VMPFC ROIs were combined into a single reward hypothesis-motivated mask for SVC. We also explored value memory responses in the MTL driven by a separate hypothesis that the hippocampus may support memory for value given its role in episodic memory. The memory hypothesis-motivated MTL ROI (including hippocampus and parahippocampal cortex) was adapted from the AAL atlas. All voxel locations are reported in MNI coordinates, and results are displayed overlaid on the average of all participants’ normalized high-resolution structural images.

**Multivariate fMRI analyses**. For multivariate classification analyses, we estimated a mass-univariate GLM where each trial was modeled with a single regressor, giving 68 regressors for the learning phase and 68 regressors for the test phase. The learning phase regressor duration modeled the 3 s anticipation period. The test phase regressor duration modeled the 4 s object re-presentation period. Models included the 6 motion regressors and block regressors as effects of no interest.

Multivariate analyses were conducting using The Decoding Toolbox (Hebart et al., 2014). Classification utilized a L2-norm learning support vector machine (LIBSVM; Chang and Lin, 2011) with a fixed cost of *c* = 1. The classifier was trained on the full reward learning phase data. The trained classifier was then tested on the full test phase data. Note that for the primary across-phase classification analysis, no cross-validation is necessary for training because no inferences are drawn and no results are reported on the training (learning phase) data.

For secondary MVPA analyses on the learning phase alone, leave-one-run-out cross-validation was used, with learning phase data subdivided into 4 blocks to increase the number of folds in cross-validation. Learning phase results were conducted across the whole brain, to avoid biasing the classification results by the univariate reward-responsive mask employed in the cross-phase classification. Also, to illustrate the SVM pattern supporting classification of high vs. low reward trials in the incidental learning phase, we trained a classifier on all learning phase data. Individual voxel weights, which represent a combination of signal and class-independent covariance, were reconstructed according to the method of (Haufe et al., 2014). After a smoothing of 6mm FWHM, the across-subject effects are depicted for illustration only.

Our primary analyses report results using the area under the receiver-operating-characteristic curve (AUC), which uses graded decision values and better accounts for biases in classification that may arise due to the different processes engaged by the incidental learning phase vs. test phase tasks. Percent correct classification is reported for the secondary analysis using incidental learning phase classification performance to filter test phase performance, as binary correct vs. incorrect learning phase classification is necessary for this procedure.

In order to constrain the classification to regions responsive to reward, we trained and tested the classifier within regions showing group-level activation to reward anticipation and outcome in the incidental learning phase. A model including the full learning phase trial showed qualitatively similar decoding accuracy. This analysis was conducted independent of feedback (“hit”) to maximize the number of trials in the analysis, as behavioral value memory performance (Experiments 1-3) did not differ due to feedback. Conducting the analyses on the subset of hit trials yielded qualitatively similar results. Test phase classification included all levels of reward value memory confidence in order to include the largest number of trials the fMRI analysis. Finally, we performed correlation analyses to examine whether behavioral performance was correlated with classification accuracy; statistical comparison of correlations were computed using Steiger’s test for differences in dependent correlations.

In a control analysis, we trained the classifier on reward during learning and tested the classifier on participant’s behavioral response during the test phase across all test phase trials and separately for correct and incorrect test phase behavioral responses. In a second control analysis, we trained the classifier on reward during learning and tested the classifier on whether test phase classifier decision values were better explained by reward associations or memory familiarity strength to confirm that any reward-related classification was not due to differences in memory.

While the above MVPA analyses were already conducted within reward-responsive regions from the incidental learning phase, we conducted a searchlight analysis for further localization using The Decoding Toolbox (Hebart et al., 2014). We used a 4-voxel radius spherical searchlight. Training of the classifier and testing were conducted as described above for the region of interest MVPA. Individual subject classification accuracy maps were smoothed with a 6mm FWHM kernel prior to group-level analysis.

## Results

### Behavioral

The experiments consisted of two phases, an incidental learning phase and a value memory test phase. We first verified that the experience of reward anticipation elicited subjective affective responses in participants. During the initial incidental learning phase, on each trial participants viewed an incidental object picture with a vertical or horizontal shaded bar indicating the potential reward value if they responded rapidly to a subsequent target (Fig 1*A*).

Our primary question was whether single affective episodes are able to support later decision making. When making a choice between two aversive options that have been only experienced once before (e.g. Kahneman et al., 1993), a value-based decision is likely to be based on the same information as a source memory judgment for the value of the past episodes. Thus, reward value-based decision making and source memory for contextual information may be two complementary ways of looking at these kind of decisions. The primary goal of our experiments was to test whether participants could remember and use the value of past episodes. As such, we explicitly instructed participants to try to recall the value of episodes (a source memory judgment) in Experiments 1-2 and in the fMRI experiment. In Experiment 3 we tested whether participant’s preference to re-engage with objects in an additional reward game – without any instruction to recall previous episodes of experience – was also supported by episodic reward experiences. Finally, in Experiment 4 we examined whether priming reward-associated experiences influenced subsequent unrelated risk-taking decisions.

### Single reward episodes support decision based on source memory for reward

In Experiment 1, after the incidental learning phase we presented participants with choices between two objects: one that had been incidentally associated with high reward during the first phase of the task and one that had been incidentally associated with low reward (Fig. 1*B*). The participants’ goal was to choose the object that had been associated with the potential for high reward, independent of feedback – i.e. reward source memory – with an incentive of €0.50 euros for correct choices. We found that choice accuracy was significantly higher than chance (65.3 ± 2.7% correct; t_(30)_ = 5.73, p < 0.0001; Fig. 2*A*, left), demonstrating that source memory decisions about value can indeed be guided by single rewarding episodes. Participants were also able to accurately assess the strength of their reward source memory, as higher confidence was associated with better performance (86.8 ± 4.9% correct at the highest confidence level).

We found no effect of successful target response (“hit”) on choice accuracy (regression coef. = 0.11 ± 0.14; t_(29)_ = 0.76, p = 0.45). The lack of an influence of feedback is surprising from a reinforcement learning perspective, as reward feedback should dominate the learning of the predictive value of objects. However, the dominance of reward anticipation over feedback mirrors behavioral affect ratings in previous studies which have demonstrated that participants experience strong positive arousal (excitement) during anticipation but only mild arousal in response to feedback (Knutson and Greer, 2008). High positive arousal during anticipation may thus dominate memory for the value of the episode, at least when only one hit or miss feedback event is associated with an object. Further, these results indicate that participants were able to follow the task instructions to choose the object that had been associated with the high reward cue information.

We also investigated whether trial-by-trial reward expectations influenced reward value memory. As the incidental learning phase order of high and low reward trials was pseudo-randomized, participants may have evolving expectations about the potential value of the upcoming trial. Whether these expectations are matched may influence value memory performance: for example, a series of high reward trials may lead to a higher expected trial value, which, when compared to the appearance of a low value cue, would lead to a negative prediction error. Indeed, when expected trial value was not matched by experienced trial value (represented as a negative prediction error), we found that participants were significantly better at selecting the alternative high reward object over the low reward objects from these trials in later choices (high reward coef. = − 0.50 ± 0.31; t_(29)_ = −1.65, p = 0.10; low reward coef. = −0.60 ± 0.27; t_(29)_ = −2.20, p = 0.028). This indicates that when high expectations are violated by the appearance of a low value trial, participants are better able to remember the low value of the object from that trial and choose the alternative high reward object.

Next, we asked whether simple differences in arousal between high and low reward experiences could account for behavioral performance when choosing between episodes of different experienced value. Specifically, we attempted to minimize the degree to which arousal, and not reward value, could support memory decisions. In theory, in Experiment 1, implicit or explicit memory for the level of arousal associated with a high-reward object could have supported behavioral performance. Thus, in Experiment 2, objects were incidentally paired with high vs. low reward or with high vs. low thermal heat pain. The incidental reward learning phase was conducted as in Experiment 1, which required a rapid instrumental response. The incidental pain learning phase, in contrast, was accomplishing using a passive Pavlovian association between an object and thermal heat pain. The test phase was similar to that of Experiment 1, with the exception that the choice prompt asked “Better experience?”. For reward source memory decisions between high reward objects vs. high pain objects, we found that accuracy was significantly higher than chance (69.1 ± 2.6% correct; t_(19)_ = 7.38, p < 0.0001; Fig. 2*A*, middle). Higher confidence was associated with better performance (86.4 ± 7.8% correct at the highest confidence level). As in Experiment 1, there was no effect of successful target response (“hit”) on choice accuracy (regression coef. = −0.32 ± 0.32; t_(18)_ = −1.00, p = 0.31). The results from Experiment 2 indicate that arousal differences are unlikely to support successful performance the kind of value source memory probes used in Experiment 1 and the following fMRI experiment. One limitation of Experiment 2 is that it is difficult to precisely match arousal across motivated reward anticipation and thermal pain, as reward-related arousal induces approach-oriented motivation, while pain-related arousal induces avoidance-oriented motivation. However, the important consideration for Experiment 2 is that the overall level of arousal between high reward and high pain is more closely matched than the level of arousal between high vs. low reward anticipation.

**Figure 2.**
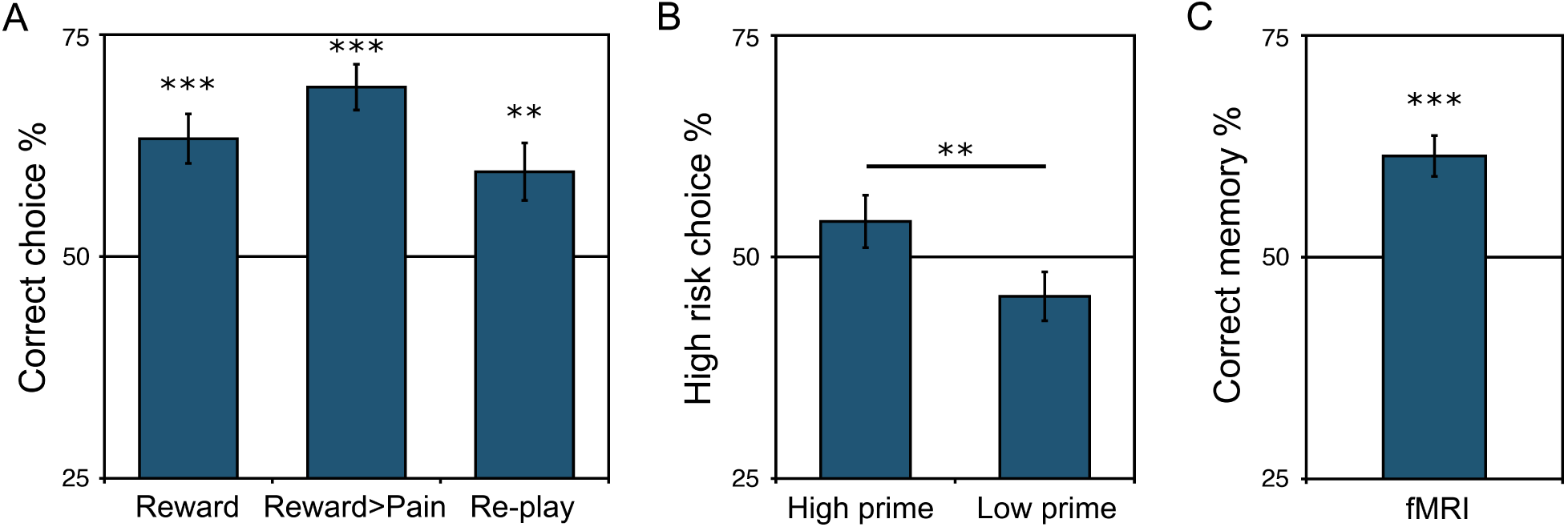
Decision making and value memory performance. ***A***, In Experiments 1-2 (“Reward” and “Reward>Pain”), accuracy in source memory – selecting the object that had been incidentally associated with high reward vs. low reward (left) or high reward vs. high pain (middle) – was significantly above chance. In Experiment 3 (“Re-play”), participants showed a significant preference to re-engage with objects that had been incidentally paired with reward (right). ***B***, In Experiment 4, objects incidentally associated with high vs. low reward exerted a significant bias on a subsequent unrelated gamble decision: participants chose the risky gamble option significantly more following objects that had been incidentally associated with high reward. ***C***, Test phase performance in the fMRI study. Participants exhibited significant source memory for the reward value of objects incidentally paired with reward in a single episode. (** p < 0.01, ***p < 0.001.)

### Single reward episodes support value-based decision making

The preceding experiments explicitly instructed participants to remember the value of the original episode during the incidental reward (or pain) phase, which relied explicitly on reward source memory. In many situations, such a source memory judgment is similar to a value-based decision based on an episode. However, recalling the value of an episode does not require participants to make a future-oriented value-based decision. Thus, in Experiment 3, we asked whether incidental reward experiences could also support decisions to re-engage with objects. In the test phase of Experiment 3, participants’ goal was to choose the object they would like to “play again” in another round of the fast response reward game (the incidental reward learning phase), allowing us to more directly test whether single reward episodes can support value-based decision making.

In the re-engagement test phase in Experiment 3, we found that participant’s decisions about which object to re-engage with were significantly directed toward the object that had been incidentally associated with high reward (59.6 ± 3.2% correct; t_(19)_ = 2.95, p = 0.0083; Fig. 2*A*, right). Higher confidence was associated with better performance (68.5 ± 6.4% correct at the highest confidence level). Similar to the previous experiments, successful target response (“hits”) did not increase performance; indeed, we observed a trending negative effect of hit on choice accuracy (regression coef. = −0.30 ± 0.37; t_(18)_ = −1.76, p = 0.08). Overall, these results demonstrate that the value of episodes has an influence beyond the effect of explicit memory for reward value (or source memory) as shown in Experiments 1 and 2, supporting the prediction that episodic reward associations can also guide future-directed decision making.

In a final behavioral study, we asked whether memory for the value of single reward experiences could also influence unrelated value-based decisions. In Experiment 4, prior to a risky gamble decision, we presented an object that had been incidentally paired with high vs. low reward in the previous incidental learning phase (Fig. 1*D*). We predicted that evoking memory for a high reward episode would activate a representation for the value of that experience and that this value may then bias participants to choose the high reward, high risk gamble option (Knutson et al., 2008). The gamble choice that followed the object picture “prime” was between a high reward €2.00 or a low reward €0.10 gamble, with even odds for winning or losing the gamble. Thus, the expected value was zero for both gambles, but the high reward gamble was associated with higher variance, or risk.

We indeed found that priming the memory of a single high reward experience increased risk-taking relative to priming the memory of a low reward experience (high reward object gamble rate, 54.0 ± 3.0%; low reward object gamble rate, 45.6 ± 2.8%; t_(19)_ = 3.48, p = 0.0025; Fig. 2***B***). Further, participants were more likely to switch from the low gamble to the high gamble following objects incidentally associated with high reward (60.0 ± 2.2%; t_(19)_ = 4.56, p < 0.001). Conversely, high reward objects were associated with a lower rate of switching to the low gamble (43.7 ± 2.1%; t_(19)_ = −3.05, p = 0.0066). As before, we found no influence of successful target response in the incidental reward phase on later behavior (high reward prime trials: coef = −0.23 ± 0.17, t_(18)_ = −1.31, p = 0.19). These results provide a novel demonstration that priming the memory of a high reward value episode increases financial risk-taking. Importantly, as the gamble was unrelated to the preceding object, participants had no motivation to strategically recall the value associated with the original experience of the object. The high reward object may have an influence on risk-taking by increasing the availability of similar positive outcome memories related to the high-risk gamble. Neurally, the influence of high reward episodes on risk-taking may be via activation of reward-related patterns of activity, as suggested by a previous report demonstrating that erotic pictures induce switches to high-risk gambles via activation in the ventral striatum / nucleus accumbens (Knutson et al., 2008).

### Reactivation of episodic value associations and source memory for reward

Building on the finding that single experiences can support value-based decisions, we conducted an fMRI study to investigate whether reactivation of value-related neural patterns could support the influence of affective episodes on memory-based decision making. To obtain neural responses to individual objects, the memory test phase in the fMRI experiment presented single objects instead of choices. On each trial, participants saw an object that had been incidentally paired with high or low reward in the preceding incidental learning phase alone for 4 s (Fig. 1*C*). After the fMRI experiment, participants rated the potential for high reward as significantly more positive than the potential for low reward (scale: 1-10; high: 7.46 ± 0.23 (mean ± SEM); low: 5.61 ± 0.35; t_(25)_ = 8.77, p < 0.001; rating data available from 26 participants), and they also rated the potential for high vs. low reward as significantly more arousing (high: 7.31 ± 0.32; low: 4.12 ± 0.30; t_(25)_ = 5.42, p < 0.001). These results verify that participants experienced high positive arousal in response to high vs. low reward anticipation.

After the incidental learning phase, participants responded with whether they thought the object had been shown with cue information signaling the potential for a high vs. low reward, a source memory judgment for a single item. Reward value memory responses were significantly above chance (61.4 ± 3.3% correct; t_(28)_ = 4.97, p < 0.001; Fig. 2*B*). Participants were also able to accurately assess the strength of their memory, as higher confidence was associated with better performance (84.9 ± 3.1% correct at the highest confidence level). Thus, incidental reward episodes can support source memory decisions for single items.

Similar to the behavioral studies, we found no effect of successful target response (“hit”) in the incidental learning phase on value memory performance (coef. = 0.13 ± 0.10; t_(27)_ = 1.34, p = 0.18), and thus subsequent fMRI analyses include all trials. Further, replicating the influence of trial value expectation on accuracy that we found in Experiment 1, we found that negative trial-value prediction errors were related to a higher rate of correct “low reward” responses for low reward trials (coef. = −0.68 ± 0.28; t_(28)_ = −2.43, p = 0.015).

A rating of object familiarity was obtained for each object after the value memory and value memory confidence responses (Fig. 1*C*). Controlling for value memory response and confidence, memory familiarity showed a trend toward being higher for objects that had been associated with the potential for high reward during incidental learning (coef. 0.11 ± 0.06; t_(25)_ = 1.81, p = 0.071). We found that familiarity ratings were related to the value memory response itself, such that familiarity ratings were higher for objects that participants had responded to with a “high” value memory response (coef. 0.25 ± 0.06; t_(25)_ = 3.95, p < 0.001). Familiarity ratings were also related to value memory confidence (coef. 0.88 ± 0.04; t_(25)_ = 24.80, p < 0.001). When including target “hit” (reward feedback) in the model, we found that feedback was negatively related to familiarity (coef. − 0.18 ± 0.06; t_(24)_ = −2.93, p = 0.0034; high reward objects only, p = 0.041; low reward objects only, p = 0.055). Although the relationship between participant’s value memory responses and confidence and memory strength demonstrates that the familiarity measure is not independent, recognition memory strength may be increased by episodic association with reward. We thus conducted control analyses for memory familiarity strength in analyses of the fMRI data. Finally, memory familiarity strength was negatively related to successfully responding to the rapid target (t_(27)_ = −2.60, p = 0.0093), suggesting that recognition was stronger for objects where a reward was missed.

We next examined our primary question of whether reward-related patterns of brain activity were significantly reactivated during re-exposure to objects in the test phase. First, we verified that univariate analyses revealed activation in expected regions to reward anticipation and feedback. High vs. low reward anticipation positively correlated with activation in regions including the left ventral striatum and medial thalamus (p < 0.05 whole-brain FWE; Table 1). Reward hit vs. miss trials also activated expected regions of interest including the ventromedial prefrontal cortex (p < 0.00001 uncorrected; Table 1) and bilateral ventral striatum (p < 0.05 FWE).

**Table 1.**
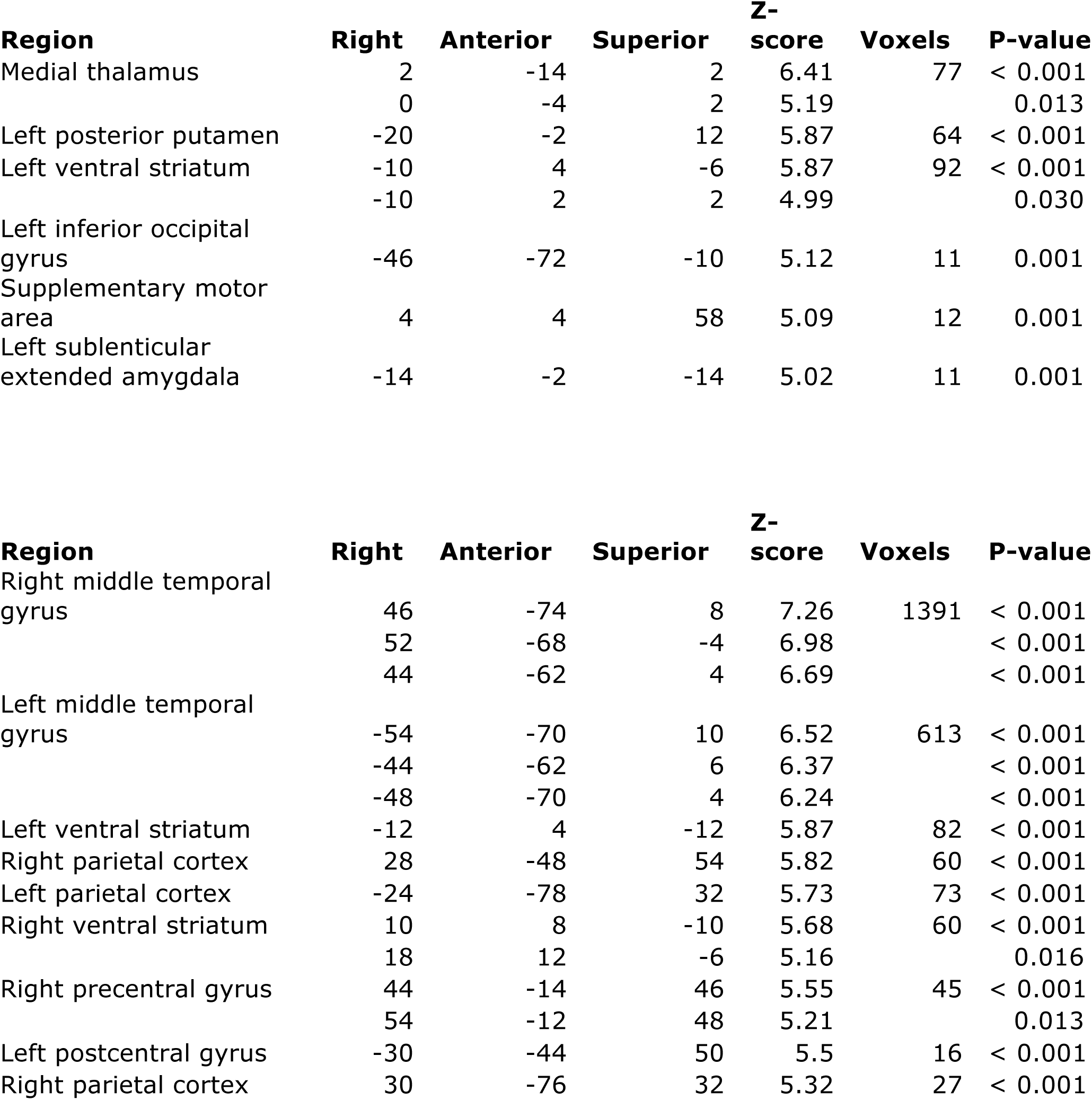
Reward fMRI experiment learning phase activation correlated with reward anticipation (top) and hit vs. miss feedback (bottom). (P < 0.05 whole-brain FWE-corrected.

For the reactivation analysis, we trained a multivoxel pattern analysis classifier on activation evoked by actual reward motivation in the incidental learning phase. A depiction of the transformed classification pattern weights discriminating high vs. low reward anticipation during the incidental learning phase, collapsing across individual patterns, is illustrated in Figure 3*A*. Qualitative inspection of the weights revealed similar effects as in the univariate contrast (Table 1), with the strongest effects in the bilateral ventral striatum, as well as the bilateral insula, anterior cingulate, thalamus, and visual cortex. We tested the performance of this classifier on activation during the test phase, when objects were presented in the absence of reward (Fig. 3*A*). This analysis allows us to detect whether distributed, multivariate value-related patterns were significantly reactivated on re-exposure to objects from the incidental learning phase.

**Figure 3.**
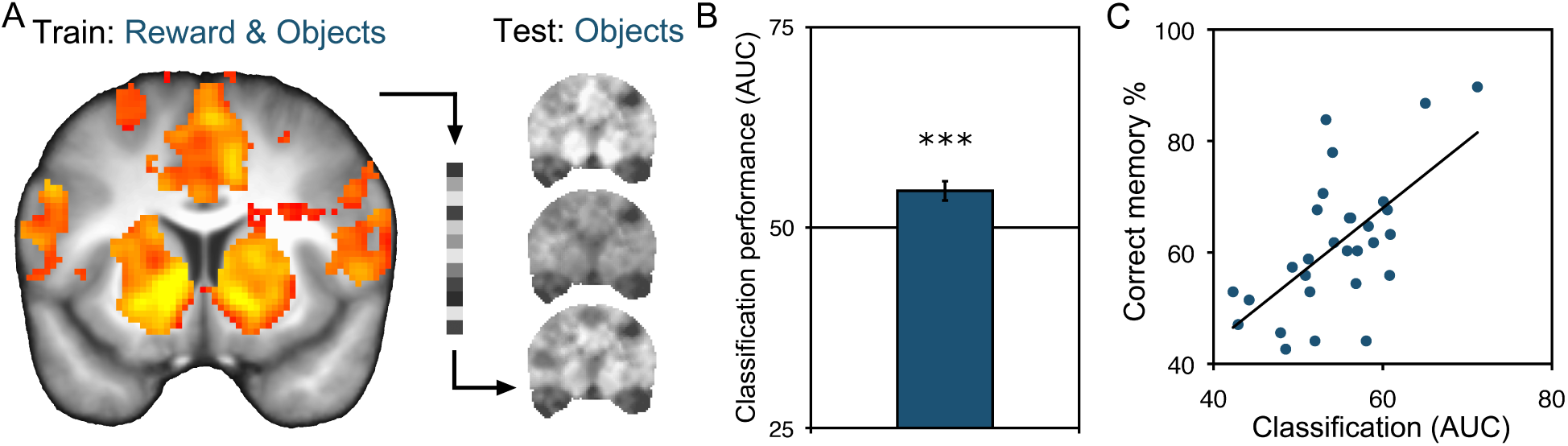
Single associations with reward lead to significant reactivation of value-related patterns during re-exposure to objects. ***A***, A pattern classifier was trained on actual reward experience and tested on re-exposure to objects in the absence of reward. Depicted is a view of the transformed support vector machine weights, collapsed across participants, illustrating the strength of positive reward classification weights in the incidental learning phase, in the reward-responsive ROI mask (Anterior = 10). **B**, Significant classification of later re-presentation of high vs. low reward objects in the test phase based on incidental learning phase patterns of activation to reward in reward-responsive regions of interest (blue). (*** p < 0.001.) ***C***, Stronger reactivation of incidental reward associations was related to better value memory behavioral performance across participants (p < 0.001).

Upon re-exposure to objects incidentally paired with reward, we indeed found significant evidence for reactivation of reward-related patterns. Within a mask defined by the main effect of reward during incidental learning, a classifier trained on reward anticipation activity in the incidental learning phase exhibited significant classification of responses in the test phase (area under the curve (AUC): 54.6 ± 1.2; t_(28)_ = 3.86 vs. chance (50.0), p < 0.001; Fig. 3*B*). Relative to the mean cross-validated reward classification performance in the incidental learning phase in the whole brain (62.5 ± 3.0 AUC; t_(28)_ = 4.19, p < 0.0001), reactivation classification was similar in magnitude to classification performance during actual reward anticipation.

Importantly, classification performance was positively related to behavioral value memory performance (r = 0.63, p < 0.001; Fig. 3*C*), such that participants with stronger patterns of reward reactivation were more likely to successfully remember the incidental episodic reward associations. The correlation was selective to test phase reactivation strength, as we found no correlation between learning phase classification accuracy and value memory performance (r = 0.01, p = 0.94). The reactivation correlation was significantly stronger (Z = 2.5, p = 0.0061), and including learning phase classification with test phase classification in a supplemental multiple regression showed a significant reactivation effect (p < 0.001) but no effect of learning phase (p = 0.70).

We tested for but did not find a difference in classification accuracy based on whether participants were correct in their individual value memory response (high reward correct vs. low: 55.6 ± 1.3 AUC; t_(28)_ = 4.42, p < 0.001; high reward incorrect vs. low: 54.0 ± 1.7 AUC; t_(28)_ = 2.32, p = 0.028; comparison, t_(28)_ = 0.86, p = 0.39). If such a difference in classification due to correct behavioral responses had been found, it would have been difficult to distinguish actual memory reactivation from test phase effects of response (high vs. low) triggering novel affective reactions. Importantly, we verified and found that the reactivation effect was not driven by a simple effect of reward memory response in the test phase: a classifier trained on reward and tested on test phase behavioral response (high vs. low) failed to show significant classification (51.85 ± 1.51 AUC; t_(28)_ = 1.22, p = 0.23). When examining correct response trials only, as expected, classification performance was significantly above chance (54.71 ± 1.60; t_(28)_ = 2.95, p = 0.0064). However, when examining incorrect response trials, the classifier failed to positively predict high vs. low response (44.27 ± 2.21; t_(28)_ = −2.59, p = 0.015). Instead, the classifier exhibited significantly below chance performance, indicating that reward-related activity at test tracked the objective reward status of the object. Finally, in an exploratory a univariate analysis of reward value memory reactivation in the test phase, we found activation in the left putamen (−32, 0, 6; Z = 4.12, p = 0.022, SVC).

We also predicted that reactivation of episodic reward experiences may differ based on how strongly participants responded to individual rewarding episodes during the incidental learning phase. We thus examined test phase classifier performance separately for those objects that had been correctly classified as being high or low reward trials in the incidental learning phase. Indeed, reactivation was only present for stimuli with stronger reward-discriminative patterns during the incidental learning phase (objects with correct learning classification, 57.4 ± 2.3%; t_(28)_ = 3.23, p = 0.0031; objects with incorrect learning classification, 49.3 ± 2.6%; t_(28)_ = −0.25, p = 0.80; comparison t_(28)_ = 1.77, p = 0.088). This result suggests that when multivariate methods are used to include only those episodic experiences with robust (or less variable) reward-discriminative patterns, re-exposure leads to stronger reactivation of reward-related patterns. In contrast, when reward-discriminative patterns in the learning phase are weaker or more variable, classification was at chance. Behaviorally, we found that performance was numerically but not significantly higher for objects with correct incidental learning phase performance (60.9 ± 2.4% vs. 58.6 ± 2.6%; t_(28)_ = 1.04, p = 0.31).

While recognition memory strength was greater for high vs. low reward items, a classifier trained on binarized memory strength success during the incidental learning phase could not predict leave-one-out classification above chance (51.2 ± 1.2%; t_(24)_ = 1.04, p = 0.31; n = 25 participants with sufficient “low” recognition strength trials). Moreover, in a multilevel regression analysis, we found that test phase reactivation (as indicated by trial-by-trial classifier decision values) was significantly predicted by the reward cue in the incidental learning phase (coef. = 0.062 ± 0.018; t_(24)_ =, p = 0.0005; across all participants) but not by graded memory familiarity strength response in the test phase (coef. = −0.002 ± 0.005; t_(24)_ = −0.40, p = 0.69; both reward and memory were entered in the same regression model).

These results demonstrate that for single episodic rewarding experiences, large-scale multivariate patterns in reward-related regions of interest during re-presentation significantly resemble those evoked by actual reward during the preceding incidental learning phase. More generally, they indicate that affect-related neural patterns are re-expressed at later recollection.

The above region of interest classification analyses demonstrated that distributed patterns of activity in reward-responsive regions show significant classification of reactivation. To further examine classification performance based on local information, we performed a searchlight analysis(Kriegeskorte et al., 2006). The searchlight analysis revealed a cluster in the left ventromedial prefrontal cortex (VMPFC; −6, 34, −10; Z = 3.32, p < 0.001 uncorrected), but this effect did not survive comparison for multiple corrections across the combined striatum and VMPFC reward-related regions of interest. The searchlight analysis also revealed a significant effect in the visual cortex (−10, −98, 0; Z = 5.87, p < 0.001 whole-brain FWE). Given the strength of the searchlight result in the visual cortex, we returned to the main classification results to examine the effect of the visual cortex on classification accuracy. While the reward-responsive mask defined from the incidental learning phase also included visual regions, excluding visual and posterior regions (significant in the searchlight analysis at a liberal threshold of p < 0.01 unc.) from the classification analysis did not qualitatively affect reactivation strength (53.6 ± 1.2 AUC; t_(28)_ = 2.86, p < 0.008).

## Discussion

We examined whether memory for the values of single experiences can guide later decision making. In our experiments, we first presented incidental, trial-unique object pictures during high or low reward episodes. Then we administered a surprise test, assessing source memory for incidental value associations or preferences to “re-play” these objects. Across three experiments, we found that participants exhibited significant memory for the value of single experiences. Moreover, in a separate experiment, we found that objects incidentally associated with reward significantly biased subsequent unrelated gambling decisions. Finally, in an fMRI experiment, we found that reward-related patterns were reactivated on re-exposure to objects. Further, individual differences in the degree of reactivation predicted value source memory behavioral performance.

As the majority of research on value-based learning has studied learning in conditions where there are many repetitions of stimulus-outcome associations (Sutton and Barto, 1998; Schultz, 2006), it has remained largely unknown whether and how the values of single episodes of experience contribute to behavior. Given the large capacity of episodic memory in humans, memory represents a rich cache of information that can support future decision making. Remembering the value of episodes may be particularly important because the circumstances associated with strongly reinforcing events are unlikely to be repeated, thus making learning via traditional gradual reinforcement learning mechanisms difficult.

In a decision making context, reactivation of the value of previous episodic experiences could be a mechanism by which episodic memory biases choice and planning (Lengyel and Dayan, 2005; Buckner, 2010). One way that memory could be useful for decision making is by providing samples of relevant past experience (Stewart et al., 2006; Weber and Johnson, 2006; Biele et al., 2009). Episodic sampling models may be able to account for aspects of learning and decision making behavior without the use of well-learned (or “cached”) values often used in reinforcement learning models (e.g. Biele et al., 2009). However, from a learning and memory systems perspective, it seems likely that both well-learned values and memory for the value of episodes influence behavior, as memory impairment due to hippocampal dysfunction does not affect the capacity of animals or humans to gradually learn the value of stimuli (e.g. Packard et al., 1989; Knowlton et al., 1996).

The broad network of regions we found to represent reward reactivation, including the striatum, VMPFC, and midbrain, are known to play an important role in learning and representing reward value (Daw and Doya, 2006; Schultz, 2006; Rangel et al., 2008). Our results suggest that in addition to supporting these processes, activity in these reward-related regions may actually represent the value of single episodes. Previous studies gave some indication of this kind of representation: for example, a recent report suggested that when participants were cued to elaborate on pre-determined specific positive life events, the VMPFC and striatum showed increased activity (Speer et al., 2014). Further, Kuhl et al. (2010) reported greater reward-related activity in the striatum and VMPFC on re-exposure, specifically when initial reward-motivated and intentionally encoded associations were successfully remembered vs. forgotten at the end of the experiment. However, with respect to the current findings, the details of the paradigm used by Kuhl et al. may be important to consider: in their experiment, a recollection memory test intervened between initial encoding and the reported fMRI effect. Successful verbal recollection of the high-reward associations in the intervening test may have been reinforcing, as it led to actual monetary rewards for the participants. Further, recent reports have shown that successful memory events during test activate reward-related regions including the striatum (Han et al., 2010; Scimeca and Badre, 2012; Schwarze et al., 2013). Thus, it is possible that reward and recollection signals during the intervening memory test contributed to later differential activation in the striatum and VMPFC for remembered vs. forgotten high-reward associations. Overall, our current results demonstrate an effect of within-participant reactivation of episodic reward associations independent of reward-motivated associative memory success.

Our fMRI results thus also provide a novel demonstration that affect-related distributed patterns of neural activity are reactivated upon re-exposure. While research in memory and emotion has demonstrated univariate overlap of mean activity between initial encoding and re-exposure (Buchanan, 2007), it has not been shown that affective patterns within the same participants are expressed at re-exposure. Our results suggest that exposure to an element of a past affective experience may lead to reactivation of similar affect-related patterns of neural activity that were expressed during the initial experience.

The hippocampus and broader medial temporal lobe did not show significant reactivation of reward value in our fMRI experiment. In contrast to this null result, previous studies have found a role for the hippocampus in representing the value of items and in imagining the value of novel experiences (Lebreton et al., 2009; Barron et al., 2013). It will be important for future studies to further explore whether the hippocampus also plays a role in representing memory for the value of episodes. Of note, the present results focus on BOLD activity during memory retrieval, and it is possible that the hippocampus plays a role in encoding episode-reward associations, similar to other sensory associations (Phillips and LeDoux, 1994; Eichenbaum and Cohen, 2001; Davachi, 2006). Indeed, previous studies have found evidence for a cooperative role of the hippocampus and reward-related regions such as the midbrain and striatum in supporting the encoding of items during states of anticipated reward as well as curiosity (Wittmann et al., 2005; Adcock et al., 2006; Wolosin et al., 2012; Gruber et al., 2014; Koster et al., 2015). Such reward-motivated encoding effects in the hippocampus have also been extended to neutral source information (Shigemune et al., 2014). Moreover, explicit goal-directed encoding of item-reward associations has been shown to engage the parahippocampus and midbrain (Dillon et al., 2014).

From a memory perspective, recalling the value of an experience is a source memory judgment, similar to memory for details or the context in which an item was studied (Cansino et al., 2002; Wheeler et al., 2006). Given the similarity between making a value-based decision from single episodes of experience and a source memory decision, our results suggest that the connection between memory and decision making research is a promising area for future studies. Interestingly, reward value is processed and encoded in an abstract manner distinct from basic sensory information (Schultz, 2006) and decisions based on value rely on different processes than perceptual decisions (Grueschow et al., 2015). Because of its unique properties, reward value can by itself support an agent’s decision making, in contrast to other sensory or memory information, to which value must be added or computed. However, previous studies have shown some overlap in univariate activation in the neural regions engaged in value-based learning and memory recollection (Foerde and Shohamy, 2011; Scimeca and Badre, 2012; Elward et al., 2015). Finally, source memory studies on emotional context share a conceptual foundation with the present experiments (e.g. Maratos et al., 2001); however, previous studies have given little attention to source memory accuracy (or decision making), with only one study reporting behavioral accuracy for episodic emotional associations for incidental stimuli (Smith et al., 2004).

While the fMRI test phase explicitly assessed source memory for value, the lack of a difference in reactivation strength based on participant’s memory accuracy suggests that reactivation of value may occur automatically and implicitly. Such an automatic effect would align with previous reports showing automatic reactivation of sensory associations (e.g. Wimmer and Shohamy, 2012). Our behavioral gambling experiment, where reward-associated objects biased unrelated gamble decisions, provides support for the influence of automatically reactivated value associations. Moreover, previous research using emotional picture primes has demonstrated a role for picture-evoked reward-related patterns of brain activity in biasing similar gamble decisions (Knutson et al., 2008). However, previous research on source memory has found evidence for different mechanisms supporting automatic vs. controlled source retrieval (Wheeler et al., 2006). Whether automatic vs. controlled retrieval of value associations relies on different mechanisms remains an open question for future research.

Interestingly, we did not find that reward pattern reactivation differed between correct and incorrect test phase behavioral responses. A strength of this null result is that our reactivation effect cannot be explained as a simple effect of test phase response, where high vs. low reward responses themselves induce a reward-like affective state. It is possible that our design was underpowered to detect reactivation differences based on behavior. It is also likely that the behavioral paradigm we used in the current study did not capture the full extend of participant’s memory for the value of episodes. Participants often have pre-existing idiosyncratic preferences for the kinds of object stimuli we used, which likely contributed a significant source of noise to the behavioral measures. Additionally, similar to how other behavioral measures such as eye-tracking have revealed traces of relational memory strength not evident in behavior (Hannula and Ranganath, 2009), future experiments that are optimized to utilize measures such as eye-tracking and reaction time have the potential to reveal more precise measures of value association memory.

In conclusion, our results provide strong support for the ability of people to use memory for the value of single experiences to guide later behavior. Further, we provide novel evidence for a value reactivation mechanism by which memory can support adaptive behavior. Understanding overactive or underactive memory formation for and reactivation of the affective elements of experiences may have implications for the treatment of mood disorders and post-traumatic stress disorder (Hamilton and Gotlib, 2008; Shin and Liberzon, 2010) as well as drug abuse (Robbins et al., 2008).

## Acknowledgments

This work was supported by ERC-2010-AdG_20100407 and DFG SFB TRR 58 and SFB 936. We thank Lea Kampermann for assistance with data collection and translation and Lara Austermann for assistance with data collection. We thank Martin Hebart and Erin Kendall Braun for helpful discussions and Georgiana Juravle for comments on a previous version of the manuscript.

